# A single-cell transposable element atlas of human cell identity

**DOI:** 10.1101/2023.12.28.573568

**Authors:** Helena Reyes-Gopar, Jez L. Marston, Bhavya Singh, Matthew Greenig, Jonah Lin, Mario A. Ostrowski, Kipchoge N. Randall, Santiago Sandoval-Motta, Nicholas Dopkins, Elsa Lawrence, Morgan M. O’Mara, Tongyi Fei, Rodrigo R. R. Duarte, Timothy R. Powell, Enrique Hernández-Lemus, Luis P. Iñiguez, Douglas F. Nixon, Matthew L. Bendall

## Abstract

Single cell RNA sequencing (scRNA-seq) is revolutionizing the study of complex biological systems. However, most sequencing studies overlook the contribution of transposable element (TE) expression to the transcriptome. In both scRNA-seq and bulk tissue RNA sequencing (RNA-seq), quantification of TE expression is challenging due to repetitive sequence content and poorly characterized TE gene models. Here, we developed a tool and analysis pipeline for Single cell Transposable Element Locus Level Analysis of scRNA Sequencing (Stellarscope) that reassigns multi-mapped reads to specific genomic loci using an expectation-maximization algorithm. Using Stellarscope, we built an atlas of TE expression in human PBMCs. We found that locus-specific TEs delineate cell types and define new cell subsets not identified by standard mRNA expression profiles. Altogether, this study provides comprehensive insights into the influence of transposable elements in human biology.

## Introduction

The classification of human cells based on cell surface markers, and more recently, RNA expression, has led to a revolution in understanding of cell function, lineage and fate^1–4^. High quality markers correlate with the characteristics and biological processes within the cell. However, these classifications have mostly been based on analyses of well characterized reference gene models (canonical genes, CG), most of which are protein coding genes^4^. A large fraction of the human genome are transposable elements (TEs), which are now appreciated to be key regulators of development and cell differentiation, and can act as promoters, enhancers, and regulators of nearby genes^5–12^. TEs play important roles in genome evolution and can have both positive and negative effects on gene regulation and genome stability. How these TEs might shape or distinguish individual cells is unknown. An understanding of TE expression at a single cell level is critical to determining the role of TEs in lineage development, cell sub-type identification and gene regulation.

Recent advances in computational biology have led to pipelines which can assess differential expression of TEs from bulk RNA-sequencing data at locus specificity^13–18^. However, there are several challenges in probing single cells for differential expression of TEs. For both bulk and single cell RNA-seq, TE gene models are underdeveloped, TE transcript abundance is low, and the repetitiveness of TEs leads to ambiguous mapping. The number of fragments sequenced in bulk samples is typically sufficient to resolve ambiguity; however, far fewer fragments are sequenced per cell in scRNA-seq. As a result, informative reads are not observed for every cell, making model-based TE quantification a technological challenge.

In this study, we developed a computational pipeline called, “Single cell Transposable Element Locus Level Analysis of scRNA Sequencing”, or “Stellarscope”. We then used Stellarscope to determine the expression of TEs in human peripheral blood mononuclear cells (PBMCs) at single cell resolution. We found that HERV and L1 transcripts can be reliably detected in single-cell RNA-seq data, and they contribute biologically relevant information to the transcriptome. We identified novel PBMC subsets using locus specific TE expression profiles compared to CGs alone.

Some TE transcripts were unique to certain cell-type transcriptomes, and contributed to cell identity. Furthermore, locus-specific HERV transcripts were distinctly expressed in differentiated hematological cell types, and could identify new cell sub-types compared to using coding genes alone.

This single-cell-resolution multi-scale analysis of the transposable element component of the human ‘dark genome’ illustrates the influence of TEs in cell identify and fate, thus establishing a novel framework for determining lineage markers derived from transposable elements, and probing the role of sequences derived from genomic dark matter in biological tissues.

## Results

### Quantification of TE expression at locus resolution in single cells with Stellarscope

Stellarscope uses four sequential stages to provides a scRNA-seq counts matrix for TE features by reassigning ambiguous (multimapping) reads to their most probable TE locus of origin. In the first stage, alignments for each read (Figure 1A-C) are intersected with the TE annotation (Figure 1D); reads with at least one alignment to a TE locus are retained for the model. For each TE-aligned read, the best alignment score for each locus it aligns to is recorded, resulting in an initial weight matrix of reads and candidate assigned features. The cell barcode (CB) of each read is compared against the user-provided list of passing barcodes (generally known as the ‘whitelist’), and both the CB and the unique molecular identifier (UMI) are stored internally.

**Figure 1.**
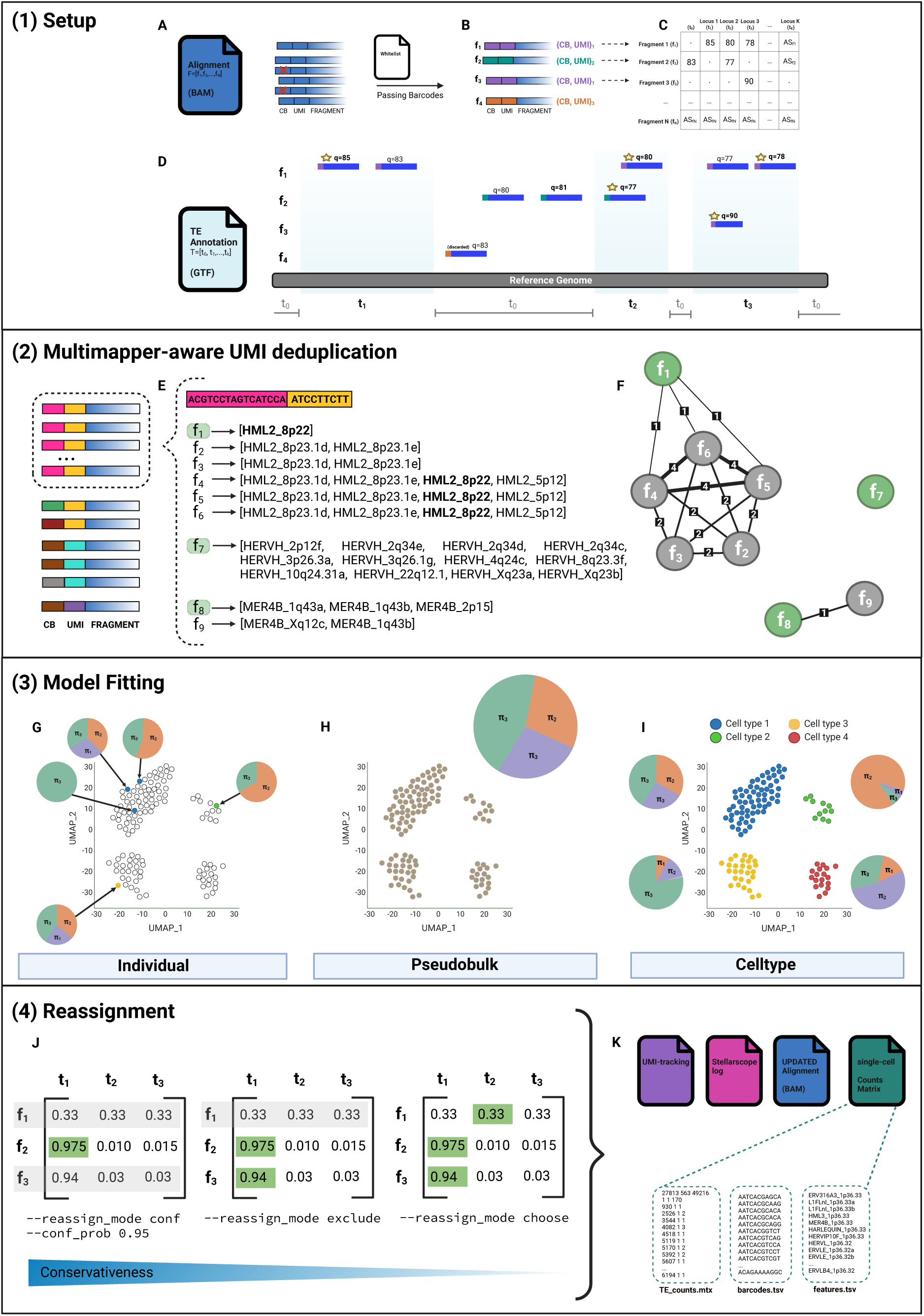
Stellarscope – Single cell Transposable Element Locus Level Analysis of scRNA Sequencing. STELLARSCOPE SETUP. (A) Alignments are filtered according to a user-provided list of passing barcodes ("whitelist"). (B) The cell barcode (CB) and unique molecular identifier (UMI) from valid fragments are stored internally. (C) Initial weight matrix with fragments as rows and candidate assigned features as columns. (D) Values for the initial weight matrix setup result from intersecting each fragment’s alignment(s) with the TE features annotation and selecting the best alignment score for each fragment for each locus. MULTIMAPPER-AWARE UMI DEDUPLICATION. (E) fragments that contain the same CB+UMI combination (i.e. duplicates) and their alignment positions are identified. (F) An undirected weighted graph is built for each CB+UMI combination with fragments as nodes and shared alignments as edge weights. For each component the most informative read according to alignment quality and ambiguity criteria is selected as representative. This method identifies and corrects non obvious duplicates (e.g. f1-f2, and f1-f3). MODEL FITTING. Stellarscope fits a Bayesian mixture model to the deduplicated weight matrix using an expectation maximization algorithm for each cell (G), for all cells (H), and for each cell type (I) in pooling modes Individual, Pseudobulk, and Celltype, respectively. REASSIGNMENT. Once the model is fitted and parameters are estimated, Stellarscope uses the posterior probability matrix to reassign ambiguous fragments to their final generating locus. Stellarscope provides a variety of reassignment strategies (J) including filtering based on a threshold, excluding fragments with multiple optimal alignments, and randomly selecting from multiple optimal alignments; these criteria result in a different number of excluded alignments (shaded in grey). The output from Stellarscope (K) includes an umi-tracking file with the graphs and representative reads selection; a log file with the fitted models, the number of observations and parameters estimated, and a log likelihood for the fitted model; an updated BAM file; and a sparse single-cell counts matrix compatible with all the generally used analysis tools.

In the second stage, PCR duplicates are identified and removed using a novel multimapper-aware UMI deduplication approach (Figure 1E). UMIs are random sequences added to DNA fragments before PCR amplification that enable identification of PCR duplicates. Sequencing fragments sharing identical UMIs are assumed to arise from the same original molecule and should only contribute one observation (count) in gene expression experiments. However, the low complexity in the UMI pool can lead to identical UMIs being attached to distinct molecules. Standard practice for UMI deduplication considers not just the UMI sequence, but also the mapping location of the sequencing fragment. This poses a problem for multimapping fragments, as the mapping location is ambiguous. Stellarscope implements an approach that considers all possible mapping locations for each read. For each UMI sequence found on multiple reads, an undirected graph is constructed with nodes corresponding to reads (Figure 1F). An edge exists between two reads if both reads have an alignment to the same locus; edges are weighted by the number of such loci. Each unconnected subgraph (connected component) represents a unique molecule, as the set of mapped genomic locations does not intersect. Reads within the same connected component are considered PCR duplicates, and the most informative duplicated read is selected as a representative. The result of this stage is a corrected weight matrix with UMI duplicates removed.

In the third stage, a Bayesian mixture model is fitted to the deduplicated weight matrix using an expectation maximization algorithm. Parameters of the model include the proportion of total reads (*π*) and the proportion of mutimapping reads (*θ*) originating from each locus. Separate models could be fitted independently for each barcoded cell, meaning that the final assignment of an ambiguous read depends solely on informative reads from the same cell (Figure 1G). In practice, this approach suffers from a lack of informative reads, due to the characteristic low expression levels of TEs and the relatively small sequencing depth per cell. To address this challenge, pooling models were implemented that enable the utilization of information across cells for resolving ambiguous reads. The “pseduobulk” pooling model estimates one set of model parameters for all cells (Figure 1H), while read membership probabilities and final assignments are determined at a single cell level. The implicit assumption of this pooling model is that the retrotranscriptome of the sample is reflective of the retrotranscriptome of each individual cell; that is, the relative expression levels of specific TE loci are similar between any two given cells. This model will perform well in samples when cellular heterogeneity is low, such as sorted cell subsets or cultured cells. In contrast, high cellular heterogeneity may lead to incorrect reassignments as TE loci that are more abundant in the sample – either due to higher expression or greater cell type proportion – will have greater weight for ambiguous reassignment. To address such cases, we implemented the “cell type” pooling model, which fits a separate model for each cell type label in the sample (Figure 1I). The cell type model assumes that the relative TE expression levels are similar among cells with the same cell type label and are not dependent on sample-level TE expression. The cell type labels are provided as input and can be determined using existing supervised or unsupervised approaches for cell type annotation.

For all three pooling modes, mixture models are specified by subsetting the initial assignments and weights for each read in the pool. Starting values for *π* and *θ*, as well as priors on these parameters, are initialized by assigning equal weight to each TE locus. The model is optimized using an expectation-maximization algorithm, which iteratively calculates read assignment probabilities and maximum a posteriori parameter estimates. The algorithm terminates when convergence is achieved or when the maximum number of iterations is reached. The outcome of this stage is the fitted models, including the read assignment posterior probabilities and estimates of *π* and *θ*. The number of observations, the number of parameters estimated, and complete data log likelihood for the fitted model are also reported, which can be used for model selection.

### Stellarscope can determine the retrotranscriptome of human peripheral blood mononuclear cells at single cell resolution

We examined the contribution of TE loci to single cell transcriptomes by profiling TE-derived transcripts in human peripheral blood mononuclear cells (PBMCs). Sequencing reads were aligned to the human genome (hg38) using alignment parameters that report up to 500 high-scoring alignments for “multimapping” reads – sequencing fragments that do not uniquely align to the reference genome (STARsolo^19^). Multimapping reads were reassigned to the most probable location using a Bayesian mixture model implemented in Stellarscope (see Methods section). UMI counts for TEs reported by Stellarscope were joined with canonical gene (CG) UMI counts for downstream analysis (Figure 2A).

**Figure 2.**
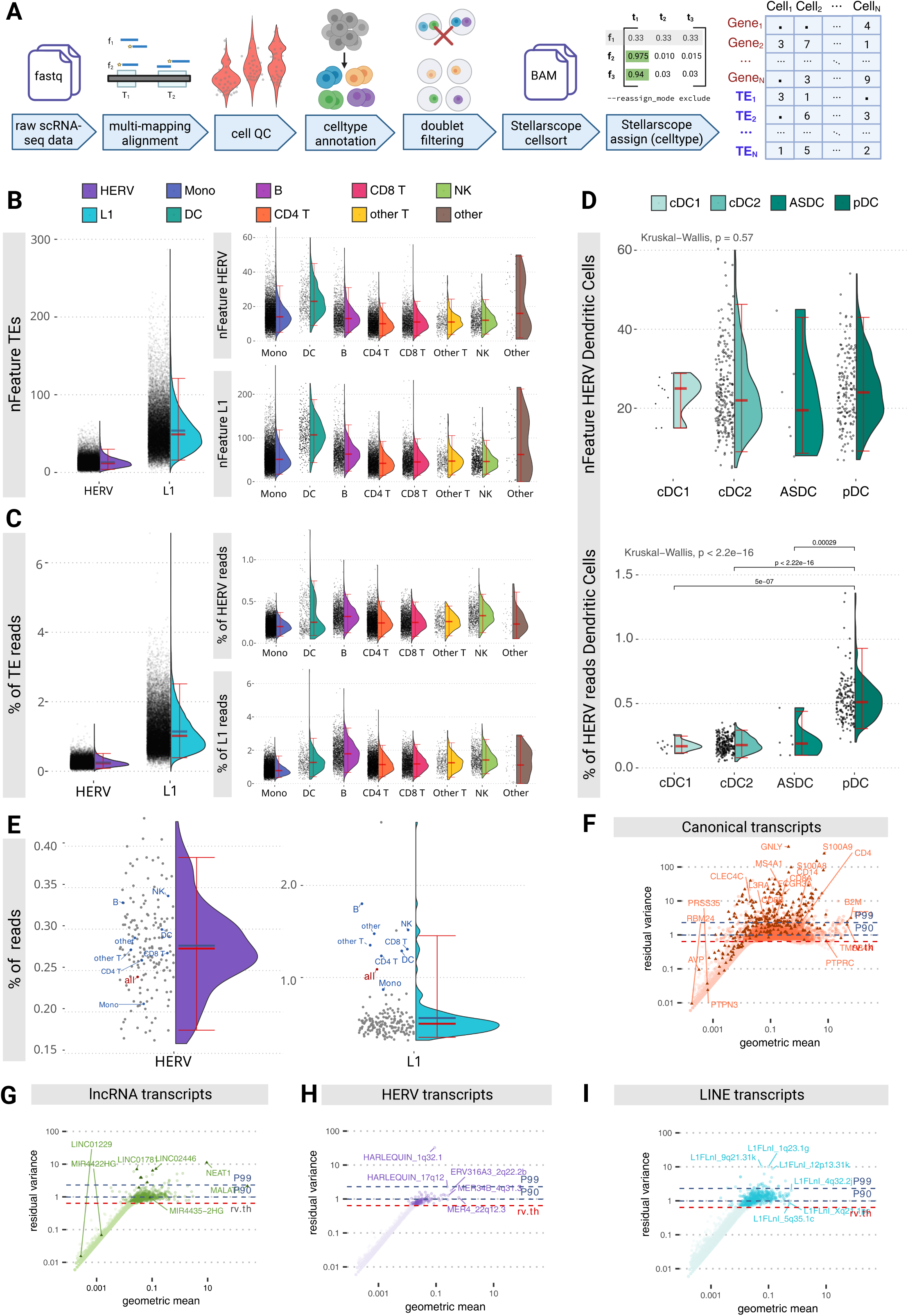
Stellarscope determines the retrotranscriptome of human PBMCs at single cell resolution. (A) Schematic representation of the stepwise analytical pipeline employed to obtain the scRNA-seq matrix counts for the PBMCs. Alignment: reads are aligned to the hg38 reference genome retaining multimappers and alignments of varying quality. Cell quality control: outlier cells are identified and excluded using adaptive thresholds on the number of features detected, total number of molecules detected, and percentage of mitochondrial reads. Celltype annotation: scRNA-seq data is projected onto the Azimuth reference atlas for PBMCs ^36^. Doublet filtering: doublets are detected and removed using Scrublet^33^ to obtain the final list of cell barcodes that are input to Stellarscope for reassignment of ambiguous reads and counting of TE features. (B) Violin plots showing the distribution of detected TE features in the PBMCs (left panel), and detected HERV features and L1 features (Y axes) by PBMC cell type (X axes). (C) Violin plots showing the distribution of the percentage of reads by cell that is contributed by TE loci in the PBMCs (left panel), the percentage of reads by cell that is contributed by HERV loci and L1 loci (Y axes) by PBMC cell type (X axes). (D) Violin plots showing comparable distributions of detected HERV features number in Dendritic Cells (upper panel) and a significant difference in the percentage of reads by cell that are contributed by HERV loci in plasmacytoid Dendritic Cells. (cDC1s: Conventional type 1 dendritic cells) (E) Measurements of TE expression from true bulk RNA-seq data for 157 PBMC samples (gray) and from pseudobulk aggregation of the PBMCs scRNA-seq data by cell type (blue) and by total cells (red). (F-I) Biological variability of expression values for different sets of transcripts by biotype showing matching patterns for TE (HERV and L1) transcripts and biologically relevant noncoding trancsripts (ncRNAs). Features with a rv value higher than 1 show relevant variation across the cells. Triangles in canonical transcripts indicate marker genes determined by the reference transcriptome.

First, we asked whether single cell expression profiling, given the low UMI counts per cell, would yield any detectable TE expression. We identified a median of 61 TE features detected per cell, with HERV and L1 features accounting for 12 and 49 features, respectively (Figure 2B). Compared with canonical genes, TEs contribute on average of 2.8% of the total features detected in each cell. The number of TE transcripts observed per cell (UMI counts) was between 2 and 655, with TEs accounting for ∼1.3% of UMI counts per cell (Figure 2C). To determine whether any PBMC cell subtypes have distinct levels of TE expression, we used cell type predictions obtained by reference mapping^20^. Dendritic cells express more TE features than other cell types, with a median of 23 HERV and 107 L1 features detected per cell (Figure 2B). However, the average proportion of TE transcripts observed was not similarly elevated in dendritic cells (Figure 2C), suggesting that expression levels at many loci is small enough that TE load is not appreciably affected. Intriguingly, we observed a bimodal distribution in the proportion of HERV transcripts for dendritic cells, indicating distinct levels of HERV expression within the same cell type. Using more specific sub-cell type labels (predicted.celltype.l2) we found that plasmacytoid dendritic cells (pDC) had significantly higher HERV loads than other dendritic cell subtypes (Figure 2D). There were no significant differences in the number of HERV features among conventional dendritic cells (cDC1, cDC2), AXL+ dendritic cells (ASDC), and pDCs (Figure 2D). Overall, we found that TE expression was detectable using single cell expression profiling, and although the contribution of the retrotranscriptome is small, it yields detectable signal that distinguishes cell types.

Second, we compared TE expression measurements obtained using bulk and single cell RNA-seq to investigate whether the different approaches would detect similar numbers of TE features and proportions of TE reads. We obtained bulk RNA-seq data from 157 PBMC samples collected from healthy donors aged 20-74. Sequencing reads were aligned using similar alignment parameters; TE expression was quantified using Telescope^13^ with identical TE annotations. Pseudobulk expression profiles were created by aggregating single cell UMI counts for the entire sample, and for each predicted cell type. We found that the proportion of HERV UMI counts (when compared to total UMI counts) in the pseudobulk dataset (0.24%) was comparable to the proportion of HERV fragments in the bulk datasets (range: 0.16%-0.43%, mean=0.28%) (Figure 2E). The proportion of L1 transcripts in the pseudobulk dataset (1.09%) was greater than nearly all bulk dataset L1 proportions (range: 0.32%-2.69%, mean=0.52%).

We hypothesize that the disparity could be attributed to the annotation quality and genomic locations of L1 loci differing from that of HERV loci. L1 annotations are more frequently located intronically or overlapping exons than HERV annotations, potentially favoring the detection of L1 transcripts by the 3’ tag-based protocol from 10x. We observed a higher number of both HERV and L1 features in single cell data (Figure S2), consistent with prior findings^21^. This may be explained by differences in sequencing depth: the pseudobulk dataset contained over 142M UMI counts, while the average size of the bulk RNA-seq datasets was less than 15M fragments; increased sequencing depth makes it more likely that low abundance transcripts will be detected.

We sought to characterize TE loci with high biological heterogeneity in the data, because these features are informative for ascribing biological characteristics to individual cells^22^. In order to separate technical variance from biological effects, we used the residual variance from models fitted to each feature to quantify how variable is their expression throughout the cells. The residual variance of most canonical (or coding) transcripts ranges between 1 and 10% (Figure 2F). TEs tend to have lower residual variance (between 1-2%) compared to canonical genes (Figures 2H and 2I). The residual variance of L1 elements was greater than the residual variance of HERVs, but for both biotypes it was in the same range as the residual variance for long-noncoding RNA transcripts (Figure 2G). There are transcripts in all biotype sets with no biological variability, including canonical transcripts annotated as marker genes, and TE features with higher residual variance than marker genes, suggesting the expression of HERVs and L1s is not merely transcriptional readthrough or random noise in RNA-seq datasets; instead, there is a deliberate regulation of a specific set of TE transcripts. Stellarscope provides information about the intricate landscape of TE expression within single PBMCs. Demonstrating that HERV and L1 transcripts can be reliably detected in single-cell data, we found that they contribute to the complexity of the transcriptome.

### Novel PBMC subsets are identified using locus specific TE expression profiles compared to canonical genes alone

Resolution of gene expression at the single cell level has revealed novel cell types and subsets. Since these studies were performed using established gene models that mostly exclude TEs, we asked whether HERV or L1 expression profiles contain distinct patterns that could inform novel cell classifications, or whether the previous cell type identities based on CGs were adequate for characterizing cells. Using different sets of highly variable features (HVFs) that include or exclude TEs, we performed linear dimensionality reduction using principal component analysis (PCA). Significant PCs were transformed using non-linear Uniform Manifold Approximation and Projection (UMAP) for visualization. Using the complete set of HVFs (including CG, HERV, and L1) yields a representation that clearly distinguishes major PBMC lineages and cell types (Monocytes, Dendritic, B cells, T cells, NK cells), as well as many cell type subsets (Figure 3A), which will help to elucidate the mechanisms underlying observed associations of dysregulated TE expression with autoimmunity, neurodegeneration, and cancer. In order to better understand the contribution of TEs to the cellular transcriptional landscape, we next performed dimensionality reduction on sets of HVFs partitioned by feature class. Dimensionality reduction using CGs alone revealed a representation that is similar to the full HVF set (Figure 3B). This was expected, as CGs include 10,982 features, over 93% of HVFs, and include HVFs with the greatest biological variability. Differences between these projections indicate information contributed by TEs.

**Figure 3:**
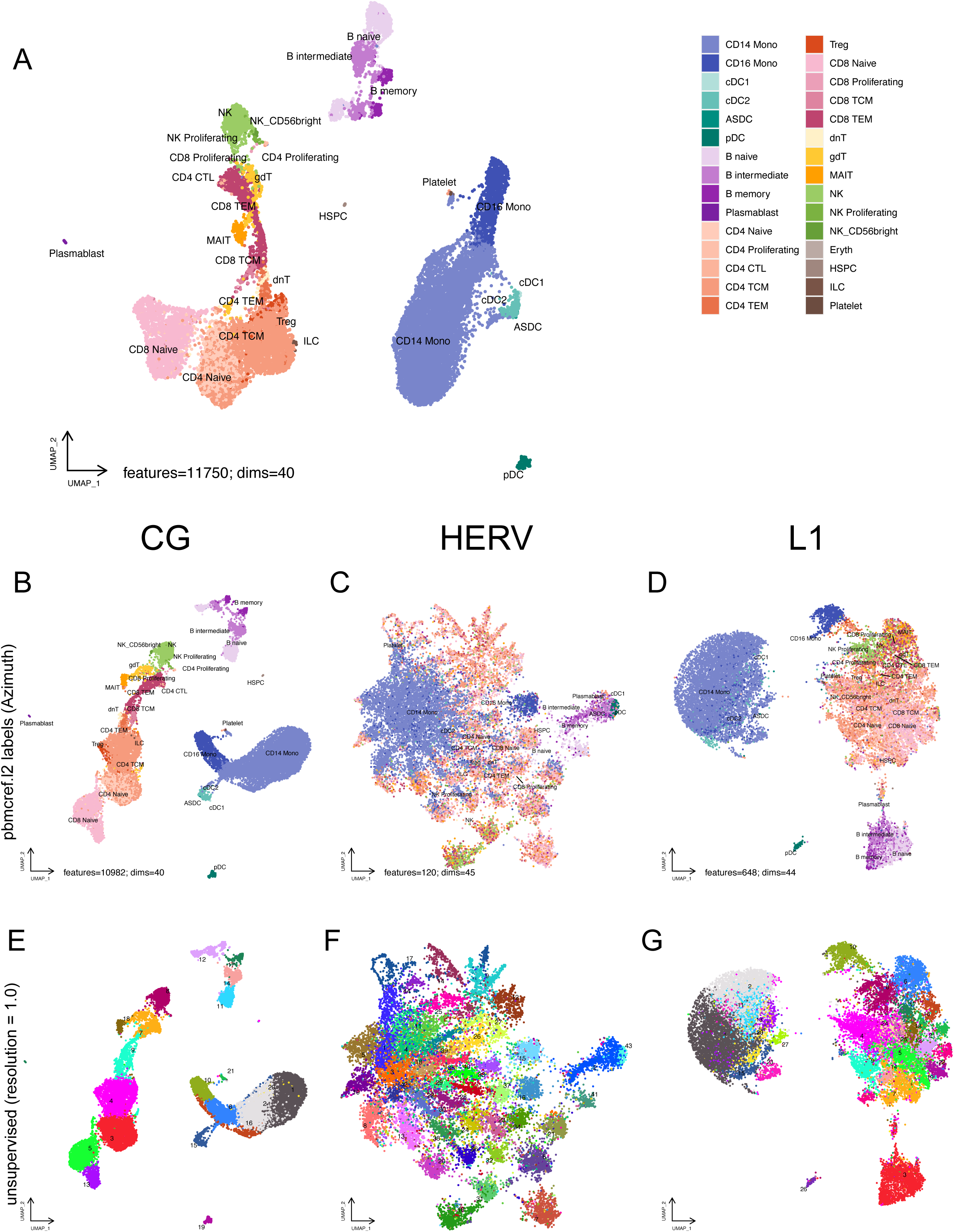
Transposable element features inform novel cell relationships and subtypes. (A) UMAP based on all identified highly variable features (HVFs), including canonical genes (CG), HERV, and L1. Cells are colored according to predicted celltype based on reference mapping to the HuBMAP human PBMC reference, celltype.l2 annotation, using Azimuth. UMAP is based on the top 40 principal components for 11,750 highly variable features. (B) UMAP based only on CG HVFs. Cells are colored as in (A), based on 40 principal components for 10,982 highly variable features. (C) UMAP based only on HERV HVFs. Cells are colored as in (A), based on 45 principal components for 120 highly variable features. (D) UMAP based only on L1 HVFs. Cells are colored as in (A), based on 44 principal components for 648 highly variable features. (E) UMAP based only on CG HVFs, as in (B). Cells are colored according to unsupervised cluster label using resolution = 1. (F) UMAP based only on HERV HVFs, as in (C). Cells are colored according to unsupervised cluster label using resolution = 1. (G) UMAP based only on L1 HVFs, as in (D). Cells are colored according to unsupervised cluster label using resolution = 1.

Projections based only on HERV HVFs were distinct from the full HVF set and describe distinct similarity patterns among cells (Figure 3C). The HERV-based projection shows some distinctions between major PBMC cell types, with separate groupings for CD14 monocytes, CD16 monocytes, B cells, Dendritic cells, and NK cells. However, some cell type subsets were not clearly distinguished. For example, there was no clear separation between CD4+ and CD8+ T cells. Furthermore, the groupings appeared *noisy* when visualized with reference-based cell type assignments. For example, although most CD14 monocytes appeared together on the left side of the UMAP, there were also CD14 monocytes in nearby groupings primarily comprised of T cells. Despite this failure to recapitulate CG-based identities at the subset level, there appears to be structure in the HERV expression patterns driving similarities among cells, in contrast to random noise. The small number of HVF HERVs (120 features) and the relatively low biological heterogeneity of these features certainly contribute to these differences, but it may also reflect novel cell states or processes involving HERV that are distinct from established celltype identities.

Similarly, LINE-1 only transcriptomes more distinctly reproduced the separation of PBMC subtypes when compared to LTR-only transcriptomes and utilized 648 features and 44 dimensions (Figure 3D).

A key hypothesis tested by this study was the potential for the addition of the retrotranscriptome to determine previously unidentified subcategories of cell types from scRNA-seq tools. When utilizing unsupervised clustering algorithms on canonical CG (Figure 3E), LTR-only (Figure 3F) at resolution = 1.0, a number of subclusters are determined distinct from the reference annotations. Where unsupervised clustering of CG-only and LINE-1 transcriptomes largely recapitulates the reference clustering within the same UMAP space, unsupervised clustering of LTR-only transcriptomes identified expression similarities as subclusters within broader cell types and these subclusters include cells from a number of cell types such as NK cells and CD4+ T cells, yet B cells and pDCs are still distinct as LTR-only clusters.

This result provides evidence that TE transcripts are unique to certain cell-type transcriptomes and contribute to cell identity. These findings are a compelling argument for the inclusion of the non-canonical TE transcriptome in analyzing scRNA-seq data and cell-type transcriptional profiles in healthy and disease conditions.

### PBMC subtypes are characterized by expression of specific HERV loci

Groups of similar cells are typically classified using markers, including surface proteins and more recently, RNA expression. High quality markers correlate with the characteristics and biological processes within the cell. Most of the markers commonly used are protein coding genes, while several long non-coding RNAs (lncRNAs) have been found to be highly sensitive in transcriptomic studies^23^. The utility of TE-derived RNAs as markers has not previously been demonstrated, partially due to technological challenges with assaying the expression of specific TE loci in either sorted bulk samples or single cells. Using Stellarscope, we achieved single locus resolution of TE expression in individual PBMC, and show that locus-specific HERV transcripts are distinctly expressed in differentiated hematological cell types (Figure 4). Transcriptional differences amongst cells correlated with known cell types, including subtypes within T cells, B cells and monocyte lineages. Overall, we identified 66 significant tests representing 34 distinct HERV loci with significant differences in expression in one or more cell subsets when compared with all other cells (adjusted p-value < 0.05, average log_2_ fold change > 0.25) (Figures 4A and S4).

**Figure 4.**
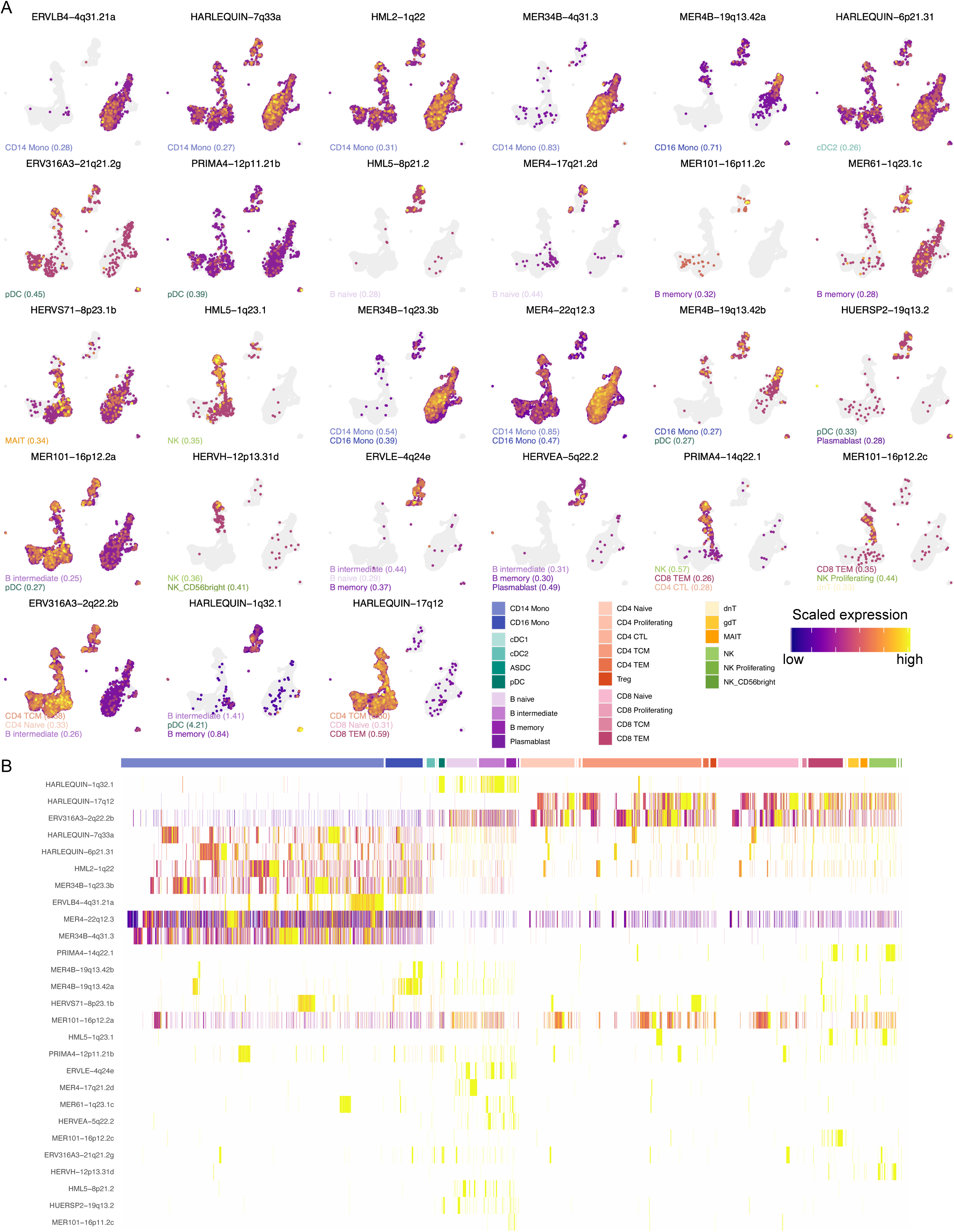
Expression of locus specific HERV features characterizes PBMC subsets. (A) Feature plots showing the relative expression in each cell for 27 HERV features with significant differential expression in one or more cell subset comparisons. Each plot is titled with the feature name; within each plot, every cell is colored according to the scaled HERV expression detected in that cell, see legend. Cells where there was no detection are colored gray. The position of cells is identical in all plots and in Fig. 3A: the cells are plotted in UMAP space calculated using all highly variable features (CG+HERV+ L1). The identity of the cell subset (or subsets) in which each HERV is significantly upgregulated is annotated in the lower left of each plot with the average log2 fold change in parentheses; HERV features that were significantly upregulated in more than three cell subsets, only the top three significance tests are shown, as ranked by adjusted p-value. Annotation text is colored according to the celltype color palettes in the legend. (B) Two dimensional heatmap showing the relative expression in each cell for 27 HERV features with significant differential expression in one or more cell subset comparisons. Each row represents a HERV feature with the names along the Y axis. Each column represents one cell with the predicted cell subset shown above the plot, colored according to the celltype color palette. Features are ordered by hierarchical clustering of the heatmap data. Cells are first grouped according to cell subset, then ordered by hierarchical clustering within each subset.

Eight HERV features were significantly upregulated in monocytes, representing possible HERV-based markers. Four features were uniquely upregulated in CD14 Monocytes, which constitute the largest proportion of cells: ERVLB4-4q31.21a (intergenic), HARLEQUIN_7q33a (intergenic), HML2-1q22, and MER34B-4q21.2. One feature, MER4B-19q13.42a, was uniquely upregulated in CD16 Monocytes, while MER34B-1q23.3b and MER4-22q12.3 were significantly upregulated for both monocyte subsets, suggesting that these may be useful in distinguishing monocytes from other cell types, but less useful in resolving subsets. Significant upregulation of MER4B-19q13.42b was detected in both CD16 monocytes and pDCs.

In pDCs, the relatively high percentage of HERV transcripts (Figure 2D) was matched by a large number of differentially expressed loci. We identified six potential HERV markers: HARLEQUIN-1q32.1, ERV316A3-21q21.2g, HUERSP2-19q13.2, MER4B-19q13.42b, PRIMA4-12p11.21b, MER101-16p12.2a. Two of these, ERV316A3-21q21.2g and PRIMA4-12p11.21b, were unique to pDCs. HUERSP2-19q13.2 and MER101-16p12.2a were shared with different B cell subtypes, while HARLEQUIN-1q32.1 was significant six different subtypes, including three B cell subtypes and two other dendritic cell subtypes. We had previously identified this locus as the HERV with the greatest biological heterogeneity (as measured by residual variance, Figure 2H) thus supporting our approach for variable feature selection. Considering the strength of this marker compared with canonical marker genes, expression of HARLEQUIN-1q32.1 in pDCs has the greatest effect size and significance of all features tested (adjusted p value < 1e-122 ; average log_2_ fold change = 4.213),and is among the top markers for all cell subtypes.

cDC2 were marked by HARLEQUIN-6q21.31. pDC were marked by ERV316A3-21q21g and PRIMA4-12p11.21b. Naive B cells were marked by HML5-8p21.2 and MER4-17q21.2d. Memory B cells were marked by MER101-16p11.2c and MER61-1q23.1c. MAIT cells were marked by HERVS71-8p23.1b, and NK cells were marked by HML5-1q23.1.

13 of the marker LTR transcripts were also detected as distinct transcriptional signatures in 2-3 cell types demonstrating the possibility of shared transcriptional events in the gene expression patterns across these cell types. pDCs were shown to share 3 marker TE transcripts: CD16 Monocytes shared the MER4B-19q13.42b with CD16 Monocytes, HUERSP2-19q13.2 with Plasmablasts, and MER101-16p12.2a with B intermediate cells. Both NK cells and the NK_CD56bright subtype shared expression of HERVH-12p13.31d.

Memory and intermediate B cell subtypes also shared differential relative expression of ERVLE-4q24e with naive B cells and HERVEA-5q22.2 with plasmablasts showing common retrotranscriptomic patterns across B lineage cells. Additionally, intermediate B cells and memory B cells shared expression of HARLEQUIN-1q23.1. CD8+ TEM also shared expression of PRIMA4-14q22.1 with and NK cells and CD4+ subtypes, as well as MER101-16p12.2c with NK proliferating cells and dnT cells. CD4+ TCM, CD4+ Naive and B intermediate cells expressed ERV316A3-2q22.2b. Finally, CD4+ TCM, naive CD8+ T cells and CD8+ TEM shared expression of HARLEQUIN-17q12, which may be a T cell lineage marker LTR transcript.

## Discussion

Cell identity has become a changing landscape with new technologies. Shape shifting cells change as they squeeze through vessels or home to tissues, and cell surface markers vary in the cell’s journey. Lineage development can be determined by cell sub-types and identification of RNA-seq transcripts from single cell resolution. Yet all of these identifiers of cell identity ignore the large part of the genome composed of transposable elements (TEs).

In this study, we present a scRNA-seq-based computational pipeline for characterizing cell identity based on the expression of human endogenous retrovirus (HERV) and Long interspersed nuclear elements type 1 (LINE-1; L1) from the TEs. We demonstrate that TEs can be identified from scRNA-seq data at a locus specific level, and that TE signatures could identify new cell sub-types over canonical gene markers and suggest a new layer of complexity of cell identity.

The initial step of the reference pipeline Stellarscope is the mapping stage, where alignments are filtered according to a list of passing barcodes. Then PCR duplicates are identified and removed using a novel multimapper-aware UMI deduplication approach. Stellarscope implements an approach that considers all possible mapping locations for each read. For each UMI sequence found on multiple reads, an undirected graph is constructed with nodes corresponding to reads. Then, a Bayesian mixture model is fitted to the deduplicated weight matrix using an expectation maximization algorithm. Importantly, due to the relatively small sequencing depth of TEs per cell, pooling models were implemented that enable the utilization of information across cells for resolving ambiguous reads.

Biologically, we then probed RNA-seq datasets for the contribution of TE loci to single cell transcriptomes. Using scRNA-seq data from human peripheral blood mononuclear cells as a reference, and found that compared with canonical genes, TEs contributed on average of 2.8% of the total features detected in each cell.

We sought to characterize TE loci with high biological heterogeneity in the data, because these features are informative for ascribing biological characteristics to individual cells. In order to separate technical variance from biological effects, we used the residual variance from models fitted to each feature to quantify how variable is their expression throughout the cells. TEs tended to have lower residual variance (between 1-2) compared to canonical genes (between 1-10). There were transcripts in all biotype sets with no biological variability, including canonical transcripts annotated as marker genes, and TE features with higher residual variance than marker genes, suggesting the expression of HERVs and L1s is not merely transcriptional readthrough or random noise in RNA-seq datasets. Instead, there is a deliberate regulation of a specific set of TE transcripts.

We then asked whether HERV or L1 expression profiles contained distinct patterns that could inform novel cell classifications Using different sets of highly variable features (HVFs) that include or exclude TEs, we performed dimensionality reduction using principal component analysis (PCA) and uniform manifold approximation and projection (UMAP). Using the complete set of HVFs (including CG, HERV, and L1) yields a representation that clearly distinguishes major PBMC lineages and cell types. Unsupervised clustering using only highly variable HERVs identified expression similarities as subclusters within broader cell types, and these subclusters included cells from a number of cell types such as NK cells and CD4+ T cells. However, B cells and pDCs were still formed distinct clusters using HERV features alone.

After having identified novel cell sub-types based upon differential TE expression, we probed known annotated cell types for unique TE expression. Notably, dendritic cells expressed more TE features than other cell types, with a median of 23 HERV and 107 L1 features detected per cell. Using specific sub-cell type labels, we found that plasmacytoid dendritic cells (pDC) had significantly higher HERV loads than other dendritic cell subtypes. In pDCs, the relatively high percentage of HERV transcripts was matched by a large number of differentially expressed loci. We identified six potential HERV markers: HARLEQUIN-1q32.1, ERV316A3-21q21.2g, HUERSP2-19q13.2, MER4B-19q13.42b, PRIMA4-12p11.21b, MER101-16p12.2a. Two of these, ERV316A3-21q21.2g and PRIMA4-12p11.21b, were unique to pDCs. Thus, our single cell profiling provides novel insights into cell identities by uncovering unique TE transcripts delineating known cell sub-types.

As the known role of transposable elements in biology grows, with major contributions noted in human development, aging, neurodegenerative diseases and cancer, understanding how single cells express TEs is critical to understand their roles in biology and human diseases. Our study establishes a novel pipeline for integrated analysis of comprehensive single-cell genomics and tissue datasets and provides new knowledge and opportunities for translation of the complexities of cell identities.

### Limitations of this study

Although this study provides a powerful computational pipeline to determine differential expression of TEs from scRNA-seq data, we note a few limitations. First, this study’s primary focus on human datasets limits mechanistic manipulation in animal models. The annotation of TEs in other species is more limited, and the biological behavior of TEs in other species quite different making comparison with human data moot. Second, although a broad array of annotated TEs are included in the reference set, there are other non annotated LTRs or other TEs which are not included, and as additional human genomes are sequenced telomere to telomere, polymorphisms within the TE genes will need to be accounted for. Finally, the insights gained from the human data sets will need to be validated with specific probes, and new tools developed to mark expression of TE ORFs (including specific antibodies) and locus specific TE probes.

## Methods

### Single cell reassignment mixture model

Stellarscope implements a generative model of single cell RNA-seq that rescales alignment probabilities for independently aligned reads based on the cumulative weights of all alignments to each transcript. Fundamentally, the probability that a given alignment is the “true” alignment increases when the total supporting information for that transcript is greater. The model and notation follow from Bendall et al. 2019^13^. Each sequencing fragment is comprised of three parts that are tracked by our model: 1) *F* = [*f*_1_, *f*_2_, …, *f*_*N*_], the set of *N* observed cDNA sequences from the originating transcript; 2) the corresponding cell barcodes *B* = [*b*_1_, *b*_2_, …, *b*_*N*_], where *b*_*i*_ = *b*_*j*_ for all *i* and *j* that originate from the same cell; and 3) a Unique Molecular Identifier (UMI) *U* = [*u*_1_, *u*_2_, …, *u*_*N*_] for each template molecule. Let *C* = [*c*_1_, *c*_2_, …, *c*_*M*_], be the set of *M* cells that are included in the model. Cells are categorized a priori into subsets, or “pools”, depending on the chosen pooling mode. Let **P** = [***P***_**1**_, ***P***_**2**_, …, ***P***_***D***_] be the set of D pools, and let *P* = [*p*_1_, *p*_2_, …, *p*_*M*_], be an indicator mapping each cell to the pool to which it belongs, ∀_*i*_ *p*_*i*_ ∈ **P**. For individual pooling mode, each cell is in a separate pool (∀_*i*_ *p*_*i*_ = *c*_*i*_). For pseudobulk pooling mode, all cells are in the same pool (∀_*i*_ *p*_*i*_ = 1). For celltype pooling mode, the pool assignment for each cell is provided as input for the model. For each pool, we estimate the abundance parameter ***π_P_*** = [*π*_*P*_0__, *π*_*P*_1__, …, *π*_*P*_*K*__ ] representing the proportion of total fragments originating from each of *K* annotated transcripts. In addition, we estimate the reassignment parameter ***θ***_***P***_ = [*θ*_*P*_0__, *θ*_*P*_1__, …, *θ*_*P*_*K*__ ] representing the proportion of ambiguous fragments generated by each transcript. Thus, the probability of observing fragment *f*_*i*_ with cell barcode *b*_*i*_ is given by:

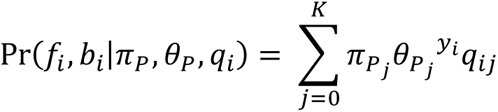

where *P* is the pool containing cell barcode *b*_*i*_ (*p*_*b*_*i*__), *π*_*P*_ and *θ*_*P*_ are pool-specific parameters, *q*_*i*_ is a vector of mapping qualities for *f*_*i*_, and *y*_*i*_ is an indicator where *y*_*i*_ = 1 if *f*_*i*_ is ambiguously aligned and *y*_*i*_ = 0 otherwise.

As in earlier work, we formulate a mixture model accounting for uncertainty in the initial fragment assignments. Let *x*_*P*_*i*__ = [*x*_*P*_*i*0__, *x*_*P*_*i*1__, …, *x*_*P*_*iK*__ ] be a set of partial assignment (or membership) weights for fragment *f*_*i*_ in pool *P*. If *f*_*i*_ did not originate from pool *P* (*p*_*b*_*i*__ ≠ *P*), then ∀_*j*_ *x*_*P*_*ij*__ = 0; otherwise 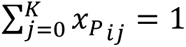 and *x*_*P*_*ij*__ = 0 if *f*_*i*_ does not align to *t*_*j*_. We assume that *x*_*P*_*i*__ is distributed according to a multinomial distribution with success probability *π*_*P*_. Intuitively, *x*_*ij*_ represents our confidence that *f*_*i*_ was generated by transcript *t*_*j*_. The complete data likelihood across all pools is

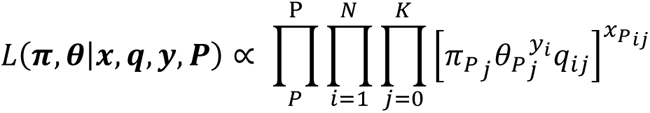

### PBMC datasets

To validate the efficacy of the Stellarscope workflow we obtained and analyzed three publicly available PBMCs scRNA-seq datasets from 10x Genomics corresponding to a healthy female donor aged 25-30. Cells were sequenced by 10x Genomics using the Chromium Next GEM Single Cell 3’ HT Reagent Kit v3.1.

### HERV annotations

A Stellarscope analysis requires an annotation that defines the transcriptional unit of each TE to be quantified. For HERV proviruses, the prototypical transcriptional unit contains an internal protein-coding region flanked by LTR regulatory regions. Existing annotations, such as those identified by RepeatMasker^24^ (using the RepBase database^25^ or Dfam^26^ identify sequence regions matching TE families but do not seek to annotate transcriptional units. Both databases represent the internal region and corresponding LTRs using separate models, and the regions identified are sometimes discontinuous. In these annotations a HERV transcriptional unit is likely to appear as a collection of nearby annotations from the same HERV subfamily.

We defined transcriptional units for HERV proviruses by combining RepeatMasker annotations belonging to the same HERV subfamily that are also located in adjacent or nearby genomic regions. Briefly, repeat families from the same HERV subfamily (internal region plus flanking LTRs) were identified using the RepBase database^25^. RepeatMasker annotations for each repeat subfamily were downloaded using the UCSC table browser^27^ and converted to GTF format, merging nearby annotations from the same repeat subfamily. Next, LTRs flanking internal regions were identified and grouped using BEDtools^28^. HERV transcriptional units containing internal regions were assembled using custom python scripts. Each putative locus was categorized according to provirus organization; loci that did not conform to expected HERV organization or conflicted with other loci were visually inspected using IGV^29^ and manually curated.

As validation, we compared our annotations to the HERV-K(HML-2) annotations published by Subramanian et al.^30^ the two annotations were concordant. Final annotations were output as GTF (S1 File); all annotations, scripts, and supporting documentation are available at https://github.com/mlbendall/telescope_annotation_db.

### Raw data/alignment

The PBMCs scRNA-seq data was publicly available and re-analyzed in this study. Raw reads were aligned to the GRCh38 reference genome using STARsolo to produce a Binary Alignment Map (BAM) file containing cell barcodes for each sequenced individual cell and for each of them, the set of possible locations their reads align to. Parameters that allow multiple alignments per read and a range score were used to retain the set of best possible alignments for each read.

### Preprocessing

Quality control was performed on the data at the cell level. Scater^31^ functions were used to identify outliers in the percentage of mitochondrial reads, total number of features, and total number of molecules detected, distributions and remove cells using these adaptive thresholds. Cell type identity was assigned to each cell using the Azimuth^32^ PBMCs reference transcriptome. Multiplets were detected using Scrublet^33^ and removed. The list of cell barcodes from cells that passed these filters was subsequently used for the Stellarscope analysis.

### Fragment Reassignment for Single-Cell Transcriptomics

The alignment from STARsolo and the list of filtered Cell Barcodes were input to Stellarscope and the BAM file alignments for valid cells were sorted using Stellarscope Cellsort. Then, Stellarscope was used in the pooling mode ‘celltype’ to reassign ambigous reads overlapping the regions from the TE annotation and obtain a TE counts matrix compatible with the Canonical Genes counts matrix.

### Downstream analysis

A merged matrix was created from the canonical genes counts matrix and the TE counts matrix and subsequent analyses were performed using Seurat version 4^34^. Cell types were annotated, and raw counts were processed using the ‘sctransform’ method to normalize the data and stabilize the variance with the aim of removing technical variability and retaining biological variability.

### Principal Component Analysis (PCA)

PCA was performed on batch-corrected Seurat data to generate a lower-dimensional representation. The data were reduced to their top 50 principal components using the “RunPCA” Seurat function.

### Clustering

Putative cell types in the three PBMC datasets were annotated by 10x Genomics. We used the 50-component PCA representation of each dataset to generate neighbour graphs and then used the Seurat function to perform the hierarchical clustering (“BuildClusterTree”).

### Differential Gene Expression

The MAST^35^ function in Seurat was applied to the single-cell PBMC datasets to identify differentially expressed genes across clusters (add parameters). MAST is a generalized linear model (GLM) framework that treats cellular detection rate as a covariate and identifies enriched genes whilst correcting for covariates and gene-gene correlations.

### Visualization

2D Uniform Manifold Approximation Projections (UMAPs) were created from the PCA matrix of the top 50 components using the “RunUMAP” function in Seurat.

### Code availability

https://github.com/nixonlab/stellarscope

## Acknowledgments

We thank Cedric Feschotte, Ethel Cesarman, Ulrike Lange, and members of their labs for many discussions about transposable elements.

