## Supplementary figures and images for "A single-cell transposable element atlas of human cell identity"

### Supplemental figure 2

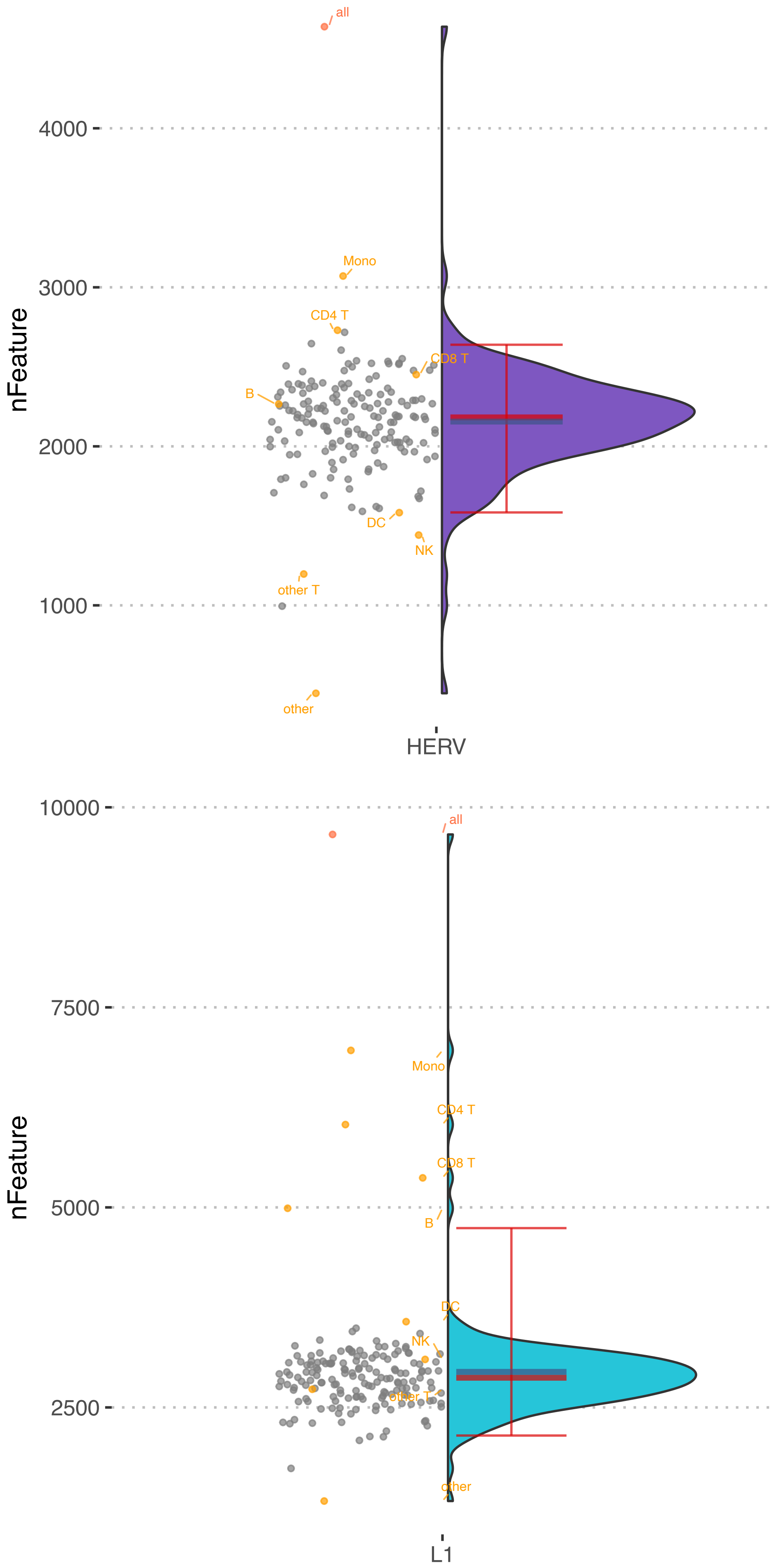
